# 50-valent inactivated rhinovirus vaccine is broadly immunogenic in rhesus macaques

**DOI:** 10.1101/053967

**Authors:** Sujin Lee, Minh Trang Nguyen, Michael G. Currier, Joe B. Jenkins, Elizabeth A. Strobert, Adriana E. Kajon, Ranjna Madan-Lala, Yury A. Bochkov, James E. Gern, Krishnendu Roy, Xiaoyan Lu, Dean D. Erdman, Paul Spearman, Martin L. Moore

## Abstract

As the predominant etiological agent of the common cold, human rhinovirus (HRV) is the leading cause of human infectious disease. Early studies showed monovalent formalin-inactivated HRV vaccine can be protective, and virus-neutralizing antibodies (nAb) correlated with protection. However, co-circulation of many HRV types discouraged further vaccine efforts. We approached this problem straightforwardly. We tested the hypothesis that increasing virus input titers in polyvalent inactivated HRV vaccine will result in broad nAb responses. Here, we show that serum nAb against many rhinovirus types can be induced by polyvalent, inactivated HRVs plus alhydrogel (alum) adjuvant. Using formulations up to 25-valent in mice and 50-valent in rhesus macaques, HRV vaccine immunogenicity was related to sufficient quantity of input antigens, and valency was not a major factor for potency or breadth of the response. We for the first time generated a vaccine capable of inducing nAb responses to numerous and diverse HRV types.

## Introduction

HRV causes respiratory illness in billions of people annually, a socioeconomic burden^1^. HRV also causes pneumonia hospitalizations in children and adults and exacerbations of asthma and chronic obstructive pulmonary disease (COPD)^2^. HRV was found to be the second leading cause of community-acquired pneumonia requiring hospitalization in US children, second only to respiratory syncytial virus, and the most common pathogen associated with pneumonia hospitalization in US adults^3^,^4^. A vaccine for HRV could alleviate serious disease in asthma and COPD, reduce pneumonia hospitalizations, and have widespread benefits for society on the whole. Decades ago, researchers identified inactivated HRV as a protective vaccine, defined virus-neutralizing antibodies (nAb) as a correlate of protection, and estimated duration of immunity^5^^−^^11^. Trials with monovalent HRV vaccine demonstrated that protection from homologous challenge and disease can be achieved with formalin-inactivated virus given intramuscularly (i.m.) or intranasally (i.n.) ^8^,^10^,^11^. Humoral immunity to heterologous virus types was not observed, though cross-reactive CD8 T cells can promote clearance^12^,^13^. Limited cross-neutralizing Abs can be induced by hyper-immunization in animals^14^,^15^. The possibility of a vaccine composed of 50, 100, or more distinct HRV antigens has been viewed as formidable or impossible^2^,^16^,^17^. There are two main challenges, generating a broad immune response and the feasibility of composing such a vaccine. The Ab repertoire is theoretically immense, and most vaccines in clinical use are thought to work via a polyclonal Ab response. Deep-sequencing of human Ab genes following vaccination against influenza virus found thousands of Ab lineages^18^,^19^. Whole pathogen and polyvalent vaccines carrying natural immunogens take advantage of this capacity. Valency has increased for pneumococcal and human papillomavirus virus vaccines in recent years. Given the significance of HRV, we tested polyvalent HRV vaccines.

There are three species of HRV, A, B, and C. Sequencing methods define 83 A types, 32 B types, and 55 C types^20^,^21^. It is thought there are 150 to 170 serologically distinct HRV types. HRV A and C are associated with asthma exacerbations and with more acute disease than HRV B^22^,^23^. HRV C was discovered in 2006 and 2007^24^^−^^27^ and recently cultured in cells^28^,^29^. Here, we focused on HRV A, the most prevalent species. There are no permissive animal challenge models of HRV virus replication, but mice and cotton rats can recapitulate aspects of HRV pathogenesis^30^,^31^. The best efficacy model is human challenge. In monovalent vaccine trials, formalin-inactivated HRV-13 was validated prior to clinical testing by assessing induction of nAb in guinea pigs, and a reciprocal serum nAb titer of 2^3^ resulting from four doses of a 1:125 dilution of the vaccine correlated with vaccine efficacy in humans^9^. Although the nAb titer required for protection is not defined, early studies established inactivated HRV as protective in humans, and immunogenicity in animals informed clinical testing.

## Results and discussion

We first used BALB/c mice to test immunogenicity. We propagated HRVs in H1-HeLa cells and inactivated infectivity using formalin. Sera from naïve mice had no detectable nAb against HRV-16 (**Fig. 1**). Alum adjuvant enhanced the nAb response induced by i.m. inactivated HRV-16 (**Fig. 1**). There was no effect of valency (comparing 1-, 3-, 5-, 7-, and 10-valent) on the nAb response induced by inactivated HRV-16 or to the 3 types in the 3-valent vaccine (HRV-16, HRV-36, and HRV-78) (**Fig. 1**). The 50% tissue culture infectious dose (TCID_50_) titers of the input viruses prior to inactivation (inactivated-TCID_50_) are provided in **Supplemental Table 1**. Original antigenic sin can occur when sequential exposure to related virus variants results in biased immunity to the type encountered first^32^. In bivalent HRV-immune mice, we observed modest original antigenic sin following prime vaccination with 10-valent inactivated HRV, and boost vaccination partially alleviated the effect (**Supplemental Fig. 1**), similar to influenza virus^32^. Collectively, these results prompted us to explore more fully the nAb response to polyvalent HRV vaccine.

**Figure 1.**
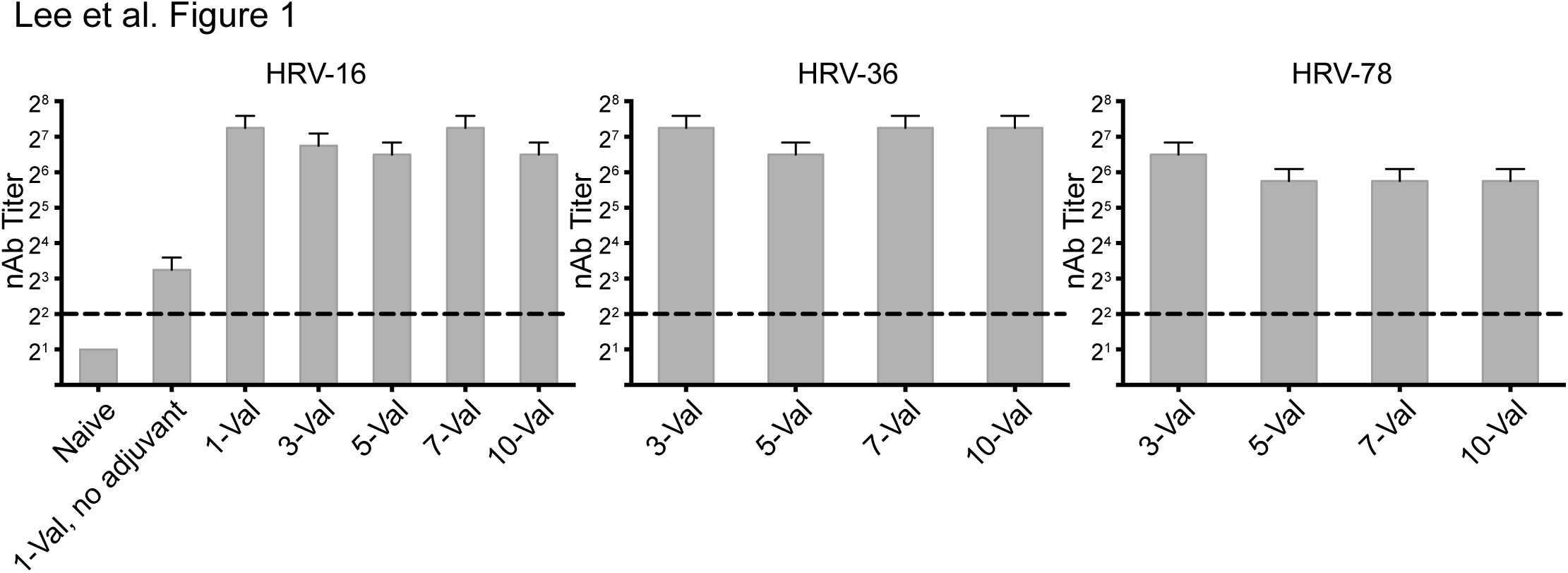
Immunogenicity of inactivated HRV is not affected by increasing valency from one to ten. Mice were vaccinated i.m. with 1-valent inactivated HRV-16 with or without alum adjuvant (5 mice per group) or with 3-valent, 5-valent, 7-valent, or 10-valent inactivated HRV with alum (20 mice per group). HRV types and inactivated-TCID_50_ doses are specified in **Supplemental Table 1**. Sera were collected 18 days after vaccination and pooled for each group. Serum nAb titers were measured against HRV-16, HRV-36, and HRV-78. The dashed line represents limit of detection (LOD). Error bars show 95% confidence interval. Data depict three independent experiments combined.

In 1975, it was reported that two different 10-valent inactivated HRV preparations induced nAb titers to only 30-40% of the input virus types in recipient subjects^33^. However, the input titers of viruses prior to inactivation ranged from 10^1.5^ to 10^5.5^ TCID_50_ per ml, and these were then diluted 10-fold to generate 10-valent 1.0 ml doses given i.m. as prime and boost with no adjuvant^33^. We hypothesized that low input antigen doses are responsible for poor nAb responses to 10-valent inactivated HRV. We reconstituted the 1975 10-valent vaccine, as closely as possible with available HRV types, over a 10^1^ to 10^5^ inactivated-TCID_50_ per vaccine dose, and we compared it to a 10-valent vaccine of the same types with input titers ranging from > 10^5^ to > 10^7^ inactivated-TCID_50_ per dose. The reconstituted 1975 vaccine resulted in no detectable nAb after prime vaccination and, following boost vaccination, nAb to the five types that had the highest input titers (**Fig. 2**). The high titer vaccine resulted in nAb to 5 of 10 types after prime vaccination and all 10 types after the boost (**Fig. 2**). Following the boost vaccinations, there appeared to be a 10^4^ inactivated-TCID_50_ per vaccine dose threshold for the induction of nAb in this model (**Fig. 2b**). Above this titer, there was no correlation between input load and nAb induction.

**Figure 2.**
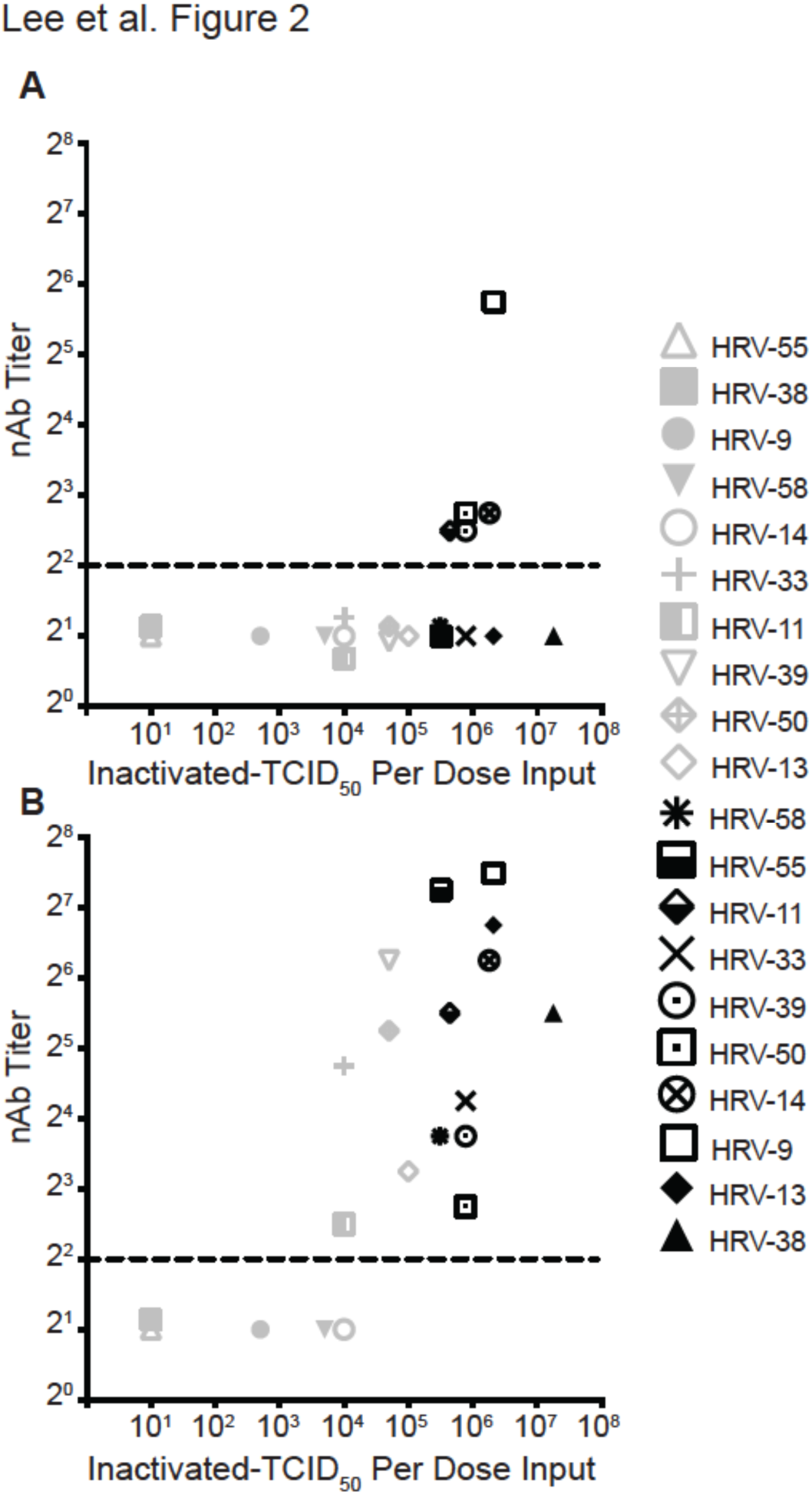
Immunogenicity of inactivated polyvalent HRV is related to dose. Mice (2 groups, 20 per group) were vaccinated with 10-valent HRV vaccine consisting of low inactivated-TCID_50_ per dose input titers (*x*-axis), similar to the 1975 Hamory et al. study^33^, plus alum (gray symbols) or with 10-valent HRV vaccine with high inactivated-TCID_50_ per dose input titers plus alum (black symbols). Sera were collected 18 days after prime (**A**) and 18 days after boost (**B**), pooled for each group, and nAb titers (*y*-axis) were measured against the indicated types in the vaccines. The dashed line represents LOD. Undetectable nAb were assigned LOD/2, and some symbols below LOD were nudged for visualization. Three independent experiments using low input titers showed similar results. There was a statistically significant association between input TCID_50_ virus titer and a detectable nAb response following prime (*P* = 0.01) and boost (*P* = 0.03) vaccination (Fisher’s exact test).

Injectable vaccines used in people are commonly given in a 0.5 ml dose. In our facility, the highest allowable i.m. vaccine volume in mice was 0.1 ml. We tested a 25-valent per 0.1 ml HRV vaccine in mice as a scalable prototype. The 25-valent inactivated HRV vaccine had a 7.4fold lower average inactivated-TCID_50_ per type per dose than the 10-valent composition (**Supplemental Table 2**) to accommodate the volume adjustment. The 10-valent inactivated HRV vaccine induced nAb to 100% of input types following the prime and the boost (**Fig. 3a)**. The nAb induced by 10-valent inactivated HRV were persisting at 203 days post-boost (**Supplemental Fig. 2**). The 25-valent inactivated HRV prime vaccination induced nAb to 18 of 25 (72%) virus types, and the 25-valent boost resulted in nAb against 24 of the 25 types (96%) (**Fig 3b**). The average nAb titer resulting from prime + boost was 2^7^ for 10-valent and 2^6.8^ for 25-valent. The data demonstrate broad neutralization of diverse HRV types with a straightforward vaccine approach.

**Figure 3.**
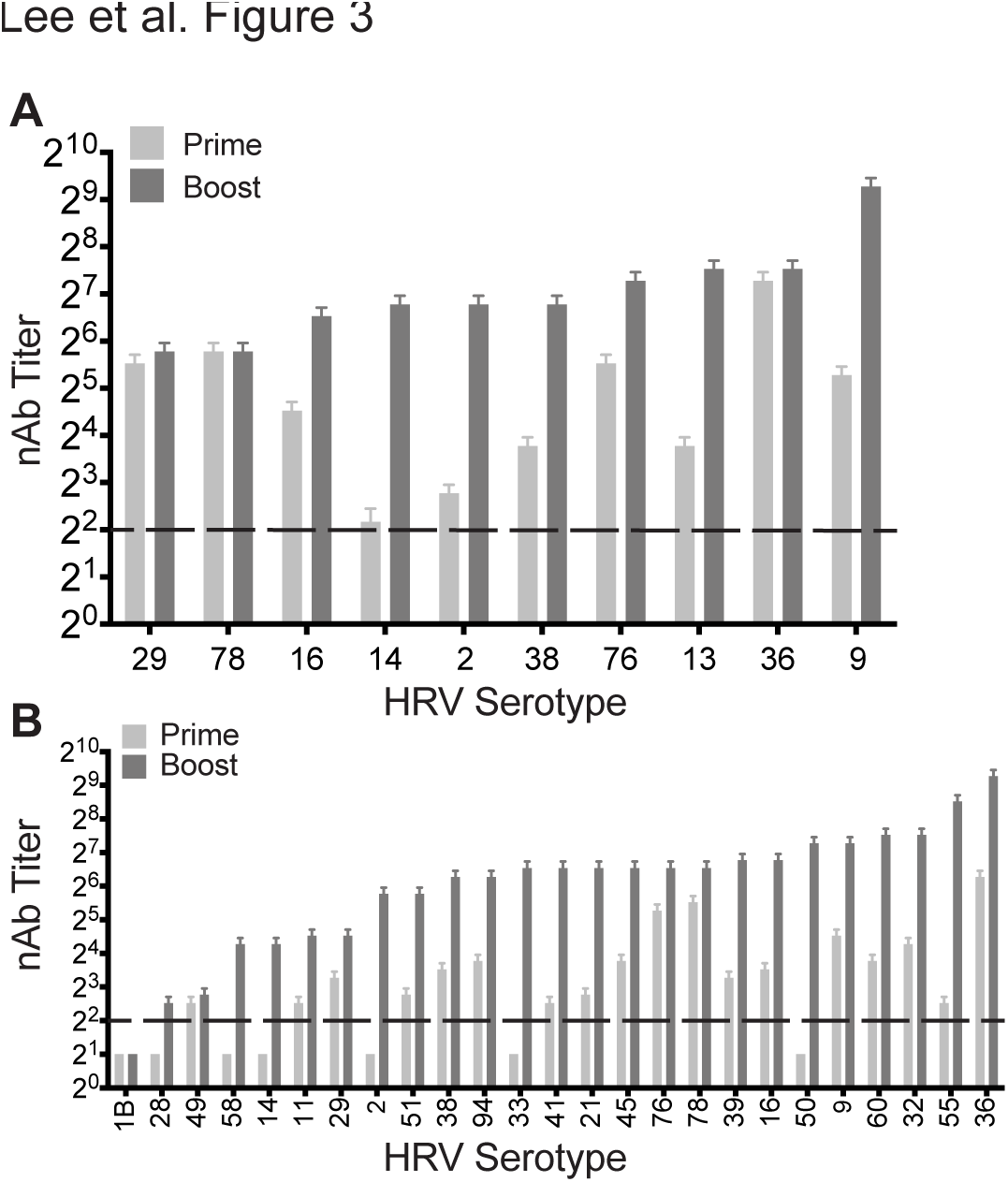
Broad nAb responses against 10-valent and 25-valent inactivated HRV in mice. The inactivated-TCID_50_ input titers per dose are specified in **Supplemental Table 2. A**, 20 mice were vaccinated then boosted at 50 days with 10-valent HRV. **B**, 30 mice were vaccinated then boosted at 50 days with 25-valent HRV. Sera were collected at day 18 (prime) and day 68 (boost). nAb levels against the indicated types in the vaccines were measured in pooled sera. Error bars depict 95% confidence interval. Data shown represent one of three (10-valent) or two (25-valent) experiments with similar results. The dashed line represents LOD. Undetectable nAb were assigned LOD/2.

In order to increase vaccine valency, we chose rhesus macaques (RMs) and a 1.0 ml i.m. vaccine volume. Two RMs were vaccinated with 25-valent inactivated HRV, and two RMs were vaccinated with 50-valent inactivated HRV. Pre-immune sera in RM A and RM B had no detectable nAb against the 25 HRV types included in the 25-valent vaccine. The inactivated-TCID_50_ titers per dose were higher in RMs than in mice (**Supplemental Table 3**). The 25-valent vaccine induced nAb to 96% (RM A) and 100% (RM B) of input viruses following the prime vaccination (**Fig. 4a**). The 50-valent vaccine induced nAb to 90% (RM C) and 82% (RM D) of input viruses following the prime vaccination (**Fig. 4c**). The breadth of nAb following prime vaccination in RM was superior to what we observed in mice, which may have been due to animal species differences and/or higher inactivated-TCID_50_ input titers in the RM vaccines. Following boost vaccination, there were serum nAb titers against 100% of the types in 25-valent HRV-vaccinated RMs (**Fig. 4b**) and 98% (49 out of 50) of the virus types in 50-valent HRV-vaccinated RMs (**Fig. 4d**). The average nAb titer resulting from prime + boost in RMs was 2^9.3^ for 25-valent and 2^8.6^ for 50-valent. The nAb responses were type-specific, not cross-neutralizing, because there were minimal nAbs induced by the 25-valent vaccine against 10 non-vaccine types (**Supplemental Fig. 3**). The nAb response to 50-valent inactivated HRV vaccine was broad and potent in RMs.

**Figure 4.**
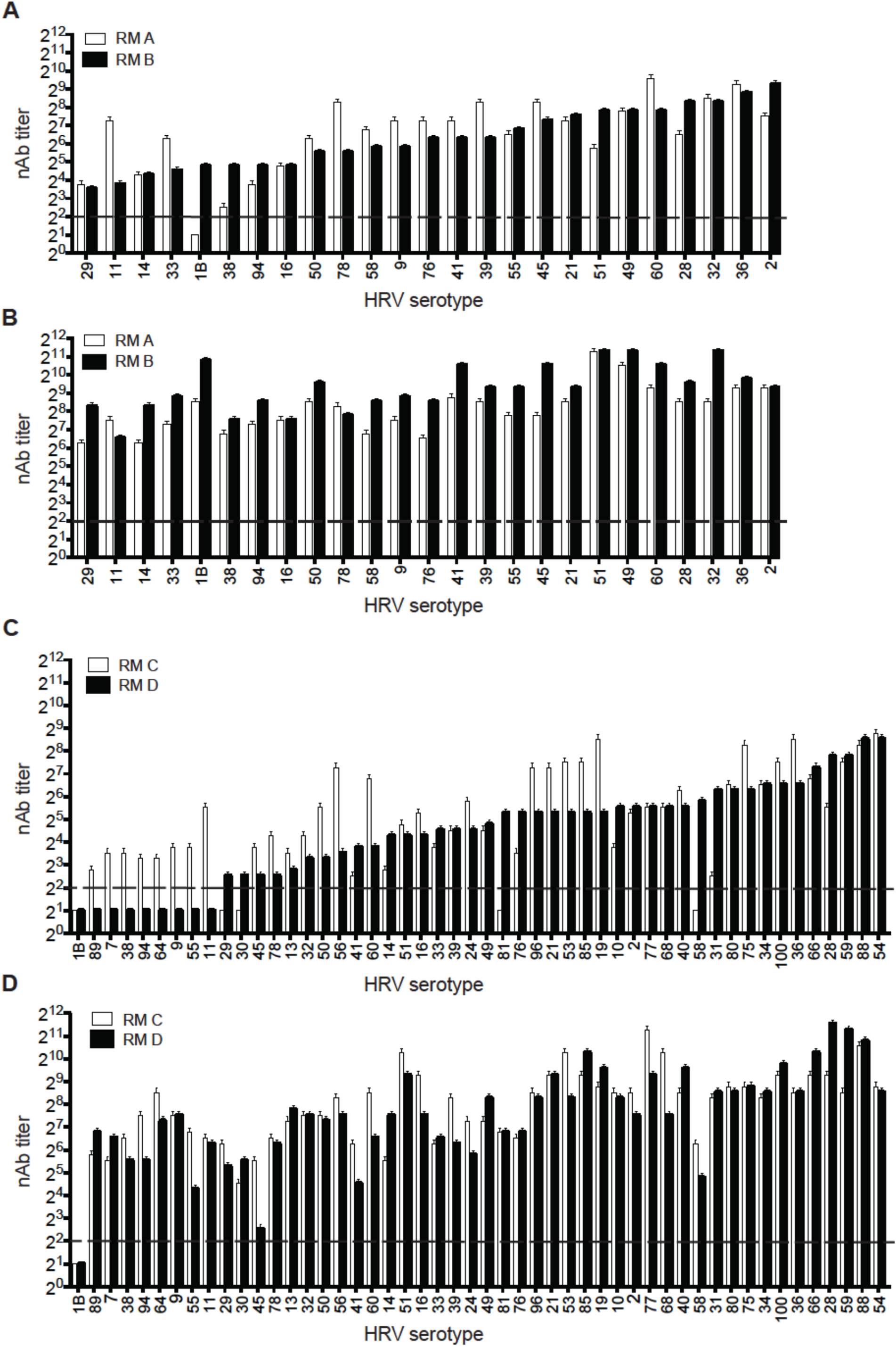
Broad nAb responses against 25-valent and 50-valent inactivated HRV in rhesus macaques. The inactivated-TCID_50_ input titers per dose are specified in **Supplemental Table 3**. Two rhesus macaques (RM A and RM B) were vaccinated i.m. with 25-valent HRV + alum (**A** and **B**), and two rhesus macaques (RM C and RM D) were vaccinated i.m. with 50-valent HRV + alum (**C** and **D**). nAb titers against input virus types were measured in individual serum samples collected at day 18 (**A** and **C**). The RM received an identical boost vaccination at day 28, and sera were collected at day 46 for determining nAb titers post-boost vaccination (**B** and **D**). Error bars depict 95% confidence interval. The dashed line represents LOD. Undetectable nAb were assigned LOD/2.

Based on our results and doses of early immunogenic HRV vaccines^6^,^8^, we estimate 10^4-5^ inactivated-TCID_50_ per type per dose will be useful. Therefore, HRV stock titers ≥ 10^7^ TCID_50_ per ml are required for a potential 83-valent HRV A formulation in a 0.5 ml dose containing alum adjuvant. The HRV stocks used in our vaccinations were produced in H1-HeLa cells, a good substrate for HRV replication but not suitable for vaccine manufacturing. We compared the infectious yield of 10 HRV types in H1-HeLa and WI-38, which can be qualified for vaccine production. Adequate yields were obtained from WI-38 cells (**Supplemental Fig. 4**). Injectable vaccines require defined purity. As proof of principle, we purified three HRV types by high performance liquid chromatography and found uncompromised immunogenicity of trivalent inactivated purified HRV in mice (**Supplemental Fig. 5**).

Forty years ago, the prospects for a polyvalent HRV vaccine were dour for good reasons^17^. However, progress in technology^34^ and advancement of more complex vaccines renders impediments to a polyvalent HRV vaccine manageable. Scale-up of HRV vaccines may be facilitated by related vaccine production processes and new cost-saving manufacturing technologies^35^^−^^37^. We provide proof of principle that broad nAb responses can be induced by vaccination with a 50-valent inactivated HRV vaccine plus alum adjuvant. Inactivated HRV has a positive history of clinical efficacy^6^,^8^,^9^. In future studies, we hope to produce a comprehensive 83-valent HRV A vaccine and generate HRV C vaccines. Our approach may lead to vaccines for rhinovirus-mediated diseases including asthma and COPD exacerbations and the common cold. Advancing valency may be applicable to vaccines for other antigenically variable pathogens.

## ONLINE METHODS

No statistical methods were used in predetermining sample sizes.

### Cell lines and viruses

H1-HeLa (CRL-1958) and WI38 (CCL-75) cells were obtained from the American Type Culture Collection (ATCC) and cultured in minimal essential media with Richter’s modification and no phenol red (MEM) (ThermoFisher) supplemented with 10 % fetal bovine serum. The cell lines were not authenticated but are not commonly misidentified (International Cell Line Authentication Committee). We tested HeLa-H1 cells using the LookOut Mycoplasma detection kit (Sigma), and these were mycoplasma negative. HRV-7 (VR-1601), HRV-9 (VR-1745), HRV-11 (VR-1567), HRV-13 (VR-286), HRV-14 (VR-284), HRV-16 (VR-283), HRV-19 (V4-1129), HRV-24 (VR-1134), HRV-29 (VR-1809), HRV-30 (VR-1140), HRV-31 (VR-1795), HRV-32 (VR-329), HRV-36 (VR-509), HRV-38 (VR-511), HRV-40 (VR-341), HRV-41 (VR-1151), HRV-49 (VR-1644), HRV-53 (VR-1163), HRV-56 (VR-1166), HRV-58 (VR-1168), HRV-59 (VR-1169), HRV-60 (VR-1473), HRV-64 (VR-1174), HRV-66 (VR-1176), HRV-68 (VR-1178), HRV-75 (VR-1185), HRV-76 (VR-1186), HRV-77 (VR-1187), HRV-78 (VR-1188), HRV-80 (VR-1190), HRV-81 (VR-1191), HRV-85 (VR-1195), HRV-88 (VR-1198), HRV-89 (VR-1199), HRV-96 (VR-1296), and HRV-100 (VR-1300) prototype strains were purchased from ATCC. HRV-1B, HRV-10, HRV-21, HRV-28, HRV-33, HRV-34, HRV-39, HRV-45, HRV-50, HRV-51, HRV-54, HRV-55, HRV-94 strains were obtained from the Centers for Disease Control and Prevention. The HRVs in the study are species A, with the exception of HRV-14 (B), and represent A species broadly^20^,^21^.

### HRV propagation and titration

HRV stocks were generated in H1-HeLa cells. Approximately 0.5 ml of HRV was inoculated onto subconfluent H1-HeLa monolayer cells in a T-182 flask. After adsorption for 1 hr at room temperature with rocking, 50 ml of HRV infection medium (MEM supplemented with 2 % FBS, 20 mM HEPES, 10 mM MgCl_2_, 1X non-essential amino acids [Gibco catalog 11140-050]) was added and the infection was allowed to proceed at 32°C in a 5% CO_2_ humidified incubator until the monolayer appeared to be completely involved with cytopathic effect (CPE), 1 to 3 days post-infection. The cells were scraped, and the cells and medium (approximately 50 ml) were transferred to two pre-chilled 50 ml conical polypropylene tubes and kept on ice while each suspension was sonicated using a Sonic Dismembrator Model 500 (Fisher Scientific) equipped with a ½-inch diameter horn disrupter and ¼-inch diameter tapered microtip secured on a ring stand. Sonication was performed by an operator in a closed room with ear protection, at 10 % amplitude, 1 sec on/1 sec off intervals, and 1 pulse per 1 ml of material. Sonication yielded higher titers than freeze-thaw. The suspension was clarified by centrifugation at 931 × *g* for 10 minutes. The supernatant was transferred to cryovials, snap-frozen in liquid nitrogen, and stored at −80°C. For comparing HRV yield in H1-HeLa and WI-38 cells, T-75 flasks of subconfluent cells were infected at a multiplicity of infection (MOI) of 0.1 TCID_50_/cell, and 20 ml of culture medium were discarded prior to scraping the cells in the remaining 5 ml followed by sonication. For all stocks, TCID_50_/ml titers were determined by infecting subconfluent H1-HeLa cells in 96-well plates with serially diluted samples, staining the cells six days post-infection with 0.1% crystal violet/20% methanol, scoring wells for CPE, and calculating the endpoint titer using the Reed and Muench method^38^.

### HRV Purification

HRV stock was harvested from H1-HeLa cell monolayers as describe above and clarified by brief centrifugation at low speed to remove large cellular debris (931 × *g*, 10 min, 4°C). In order to remove excess albumin from the crude virus stock by affinity chromatography, the supernatant was loaded onto a HiTrap Blue HP column (GE Healthcare) using an ÄKTAPurifier system (GE Healthcare) according to the manufacturer specifications. Flowthrough was subsequently loaded through a HiTrap Capto Core 700 column (GE Healthcare) to refine the virus prep by size exclusion chromatography (SEC). The flowthrough from the HiTrap Blue HP and the HiTrap Capto Core 700 was captured using the ÄKTAPurifier system (GE Healthcare) with a 20 mM sodium phosphate buffer (pH 7.0). Flowthrough from SEC was dialyzed overnight with 0.1 M Tris-HCl buffer (pH 8.0), then loaded onto a HiTrap Q XL column (GE Healthcare) and separated into fractions by ion exchange chromatography. Virus-containing fractions were eluted using the ÄKTAPurifier system (GE Healthcare) with a 0.1 M Tris-HCl buffer (pH 8.0) and a sodium chloride gradient. Fractions showing high absorption peaks at 280 nM were collected and analyzed for viral titer by TCID_50_ end-point dilution assay, and fraction purity visualized on a 10% SDS-PAGE gel by silver stain (Thermo Fisher Scientific) (**Supplemental Figure 6**). Fractions of HRV-16, HRV-36, and HRV-78 of high virus titer and purity were combined for formalin-inactivation as described below.

### Mice and Rhesus macaques

All experiments involving animals were conducted at Emory University and the Yerkes National Primate Research Center in accordance with guidelines established by the Animal Welfare Act and the NIH Guide for the Care and Use of Laboratory Animals. Animal facilities at Emory University and the Yerkes Center are fully accredited by the Association for Assessment and Accreditation of Laboratory Animal Care International (AAALAC). The Institutional Animal Care and Use Committee (IACUC) of Emory University approved these studies. Pathogen-free, 6-7-week female BALB/c mice were purchased from the Jackson Laboratory (Bar Harbor, ME, USA). Mice were randomly assigned to groups based on sequential selection from an inventory, and investigators were not blinded to outcome assessment.

Young adult (3 - 5 kg, 2 - 4 years of age, 2 females and 2 males) Indian rhesus macaques (*Macaca mulatta;* RM) were maintained according to NIH guidelines at the Yerkes National Primate Research Center. Handling and movement of RMs was performed by qualified personnel who have received specific training in safe and humane methods of animal handling at the Yerkes National Primate Research Center. The initial exclusion criterion was pre-existing nAb against HRV. The studies were conducted in strict accordance with US Department of Agriculture regulations and the recommendations in the Guide for Care and Use of Laboratory Animals of the NIH. The RMs were allocated in an un-blinded fashion to two vaccine groups (25-valent and 50-valent), one male and one female per group.

### Vaccination

Before immunization, all HRV types were inactivated by addition of 0.025% formalin followed by incubation with stirring for 72 hr at 37°C, as previously described for HRV vaccine^33^. Complete inactivation of infectivity was confirmed by end-point TCID_50_ titration in H1-HeLa cells. Formalin inactivation by this method resulted in greater immunogenicity in mice than alternative inactivation by beta-propiolactone, suggesting formalin inactivation preserved antigenic determinants. Mice were vaccinated i.m. with inactivated HRV strains mixed with 100 μg of Alhydrogel adjuvant 2% (aluminum hydroxide wet gel suspension, alum) (Sigma catalog A8222 or Invivogen catalog vac-alu) according instructions of the manufacturers. The total volume per mouse was 100 μl, administered in 50 μl per thigh. Mice were given a second identical vaccination (boost) at the time indicated in figure legends. RMs were vaccinated i.m. with inactivated HRV strains mixed with 500 μg of Alhydrogel adjuvant 2%. The total volume per RM was 1 ml, administered in one leg. RMs were boosted with an identical vaccination at four weeks.

## Serum collection

In mice, peripheral blood was collected into microcentrifuge tubes from the submandibular vein. Samples were incubated at room temperature for 20 min to clot. The tubes were centrifuged 7500 × g for 10 min to separate serum. The serum samples were pooled from mice of each group and stored at −80 °C until used. Phlebotomy involving RMs was performed under either ketamine (10 mg/kg) or Telazol (4 mg/kg) anesthesia on fasting animals. Following anesthesia with ketamine or Telazol, the animals were bled from the femoral vein. Yerkes blood collection guidelines were followed and no more than 10 ml/kg/28 days of blood was collected. After collecting blood in serum separating tube (SST), samples were incubated at room temperature for 30 min. The tubes were centrifuged 2500 × g for 15 min to separate serum. The serum samples from individual RM were stored at −80 °C until used.

### Serum neutralization assay

H1-HeLa cells were seeded in 96-well plates to attain 80-90 % confluence in 24 h. Heat-inactivated (56°C, 30 min) serum samples were 2-fold serially diluted in MEM and added to 500 TCID_50_/mL HRV of each type to be tested, in an equal volume. The virus and serum mixtures were incubated 37°C for 1 h. Then, 50 μl of the serum-virus mixture was transferred onto H1-HeLa cell monolayers in 96-well plates in triplicate, and plates were spinoculated at 2,095 × *g* for 30 min at 4°C. For each type, a no-serum control was added to test the input 500 TCID_50_. We tested pooled HRV-16 anti-sera against HRV-16 in each assay as a standard. After spinoculation, 150 μl of HRV infection medium were added to each well. The 96-well plates were incubated for 6 days at 32°C and 5% CO_2_ and then stained with crystal violet as described above. Wells were scored for the presence or absence of CPE. Neutralizing antibody endpoint titers and 95% confidence intervals were determined by the method of Reed and Muench, as previously described for HRV^14^,^38^. The 95% confidence interval indicates variability of three technical replicates within a single nAb experiment.

Supplementary Information is available in the online version of the paper.

## Acknowledgements

We are indebted to the Yerkes veterinary personnel for providing technical assistance. We thank Max Cooper (Emory University) and Joshy Jacob (Emory University) for helpful discussions. This study was supported by a pilot grant from the Emory+Children’s Center for Childhood Infections and Vaccines (CCIV) to M.L.M and in part by Department of Health and Human Services, National Institutes of Health grants 1R01AI087798 and 1U19AI095227 to M.L.M. This work is dedicated to A.R.

## Author Contributions

S.L. and M.T.N. contributed equally. S.L., M.T.N., M.G.C., J.B.J, E.A.S, A.E.K., and R.M.L. performed experiments. K.R., Y.A.B., J.E.G, and P.S. provided reagents and advice. X.L. and D.D.E. provided rhinovirus types. S.L., M.T.N., and M.L.M designed the experiments, analyzed data, and wrote the paper.

